# TDP-1 and FUST-1 co-inhibit exon inclusion and control fertility together with transcriptional regulation

**DOI:** 10.1101/2023.04.18.537345

**Authors:** Morgan Taylor, Olivia Marx, Adam Norris

## Abstract

Gene expression is a multistep, carefully controlled process, and crosstalk between regulatory layers plays an important role in coordinating gene expression. To identify functionally relevant coordination between transcriptional and post-transcriptional gene regulation, we performed a systematic reverse-genetic interaction screen in *C. elegans*. We combined RNA binding protein (RBP) and transcription factor (TF) mutants, creating over 100 RBP; TF double mutants. This screen identified a variety of unexpected double mutant phenotypes, including two strong genetic interactions between the ALS-related RBPs, *fust-1* and *tdp-1*, and the homeodomain TF *ceh-14*. Losing any one of these genes alone has no significant effect on the health of the organism. However, *fust-1; ceh-14* and *tdp-1; ceh-14* double mutants both exhibit strong temperature-sensitive fertility defects. Both double mutants exhibit defects in gonad morphology, sperm function, and oocyte function. RNA-seq analysis of double mutants identifies *ceh-14* as the main controller of transcript levels, while *fust-1* and *tdp-1* control splicing through a shared role in exon inhibition. We identify a cassette exon in the polyglutamine-repeat protein *pqn-41* which *tdp-1* inhibits. Loss of *tdp-1* causes the *pqn-41* exon to be aberrantly included, and forced skipping of this exon in *tdp-1; ceh-14* double mutants rescues fertility. Together our findings identify a novel shared physiological role for *fust-1* and *tdp-1* in promoting *C. elegans* fertility in a *ceh-14* mutant background and reveal a shared molecular function of *fust-1* and *tdp-1* in exon inhibition.

## INTRODUCTION

Eukaryotic gene expression requires coordination across multiple layers of regulatory control, including transcription, RNA processing, and translation. Two major classes of proteins responsible for this gene expression regulation are transcription factors (TFs) and RNA binding proteins (RBPs). Regulatory activities for individual TFs and RBPs have been well described, but less is known about how TFs and RBPs might coordinately control gene expression across multiple regulatory layers. Nevertheless, a growing body of recent evidence demonstrates extensive crosstalk between transcriptional and post-transcriptional factors occurs to regulate gene expression^1–5^.

RBPs regulate many aspects of RNA processing including pre-mRNA splicing, mRNA export and localization, and translation^6^. RBP dysfunction is especially notable in the nervous system, and RBP mutations have been implicated in multiple neurodegenerative diseases^7–9^. For example, the RBPs TDP-43 and FUS are involved in several RNA-related functions, including splicing and RNA transport^10,11^. Mutations in either RBP are directly linked to Amyotrophic Lateral Sclerosis (ALS) cause their mislocalization and aggregation in the cytoplasm, leading to progressive degeneration of neurons^12–15^. The disease-associated roles of RBPs such as TDP-43 and FUS have been extensively studied, but in many cases the physiological functions for these RBPs outside of disease context remain unresolved. Understanding how RBPs play a role in essential cellular functions and in the context of global gene expression coordination will be key for understanding and treating such diseases.

To identify functionally important TF-RBP gene expression coordination, we set out to systematically test for genetic interactions between TF and RBP mutants. Screens for genetic interactions, in which a phenotype occurs in a double mutant that is not predicted based on the single mutant phenotypes, have a rich history of identifying genes with related activities and/or redundant functions^16,17^. In single-celled organisms such as bacteria and yeast, genetic interaction analysis has been carried out at genome-wide scale, revealing hundreds of thousands of interactions in which a double mutant has a fitness greater than expected (positive interaction) or less than expected (negative or synthetic interaction) based on the single mutant fitness phenotypes^18,19^.

In the nematode *C. elegans*, we recently employed CRISPR/Cas9 to systematically delete evolutionarily-conserved, neuronally-expressed RBPs using homology-guided replacement to insert heterologous GFP fluorescent markers in place of the deleted gene^20^. These CRISPR/Cas9-generated RBP mutants enabled us to conduct a systematic pairwise genetic interaction screen across neuronal RBPs in *C. elegans*. We identified multiple novel synthetic interactions and revealed previously-unexplored physiological functions for several RBPs. Here, we aimed to apply this technology to investigate coordination of gene expression across regulatory layers, by conducting a pairwise genetic interaction screen between neuronally-enriched RBPs and TFs.

To do so, we generated all possible double-mutant TF-RBP combinations of 10 RBP and 11 TF gene deletion mutants, creating a total of 110 double mutants. We identified unexpected phenotypes in several double mutants, revealing extensive functional interactions between TFs and RBPs. One such synthetic phenotype was reduced fertility in *tdp-1; ceh-14* and *fust-1; ceh-14* double mutants. *tdp-1* and *fust-1* are the *C. elegans* homologs of TDP-43 and FUS, and mutations in both RBPs have been implicated in ALS^8,15,21^. Both *tdp-1; ceh-14* and *fust-1; ceh-14* double mutants exhibit reduced egg production, decreased sperm efficacy, and gonad migration defects. As *tdp-1, fust-1,* and *ceh-14* single mutants do not exhibit the same striking fertility phenotype as these double mutants, our findings identify a potential coregulatory role for these genes in gonad and sperm development. We find a shared role of *fust-1* and *tdp-1* in inhibiting exon inclusion, and identify a cassette exon in *pqn-41,* inhibited by *tdp-1*, that contributes to the fertility defects in *tdp-1; ceh-14* double mutants. Our findings thus uncover novel physiological functions for *fust-1* and *tdp-1*, in the specific context of a *ceh-14* mutant background, and shed light on their shared molecular roles.

## METHODS

### *C. elegans* strains and maintenance

All *C. elegans* strains were cultured on Nematode Growth Media (NGM) plates seeded with *E. coli.* Strains were maintained at 20°C unless otherwise stated. Some strains were provided by the CGC, which is funded by NIH Office of Research Infrastructure Programs (P40 OD010440).

Double mutant strains were created and confirmed by visualization of GFP markers and by PCR.

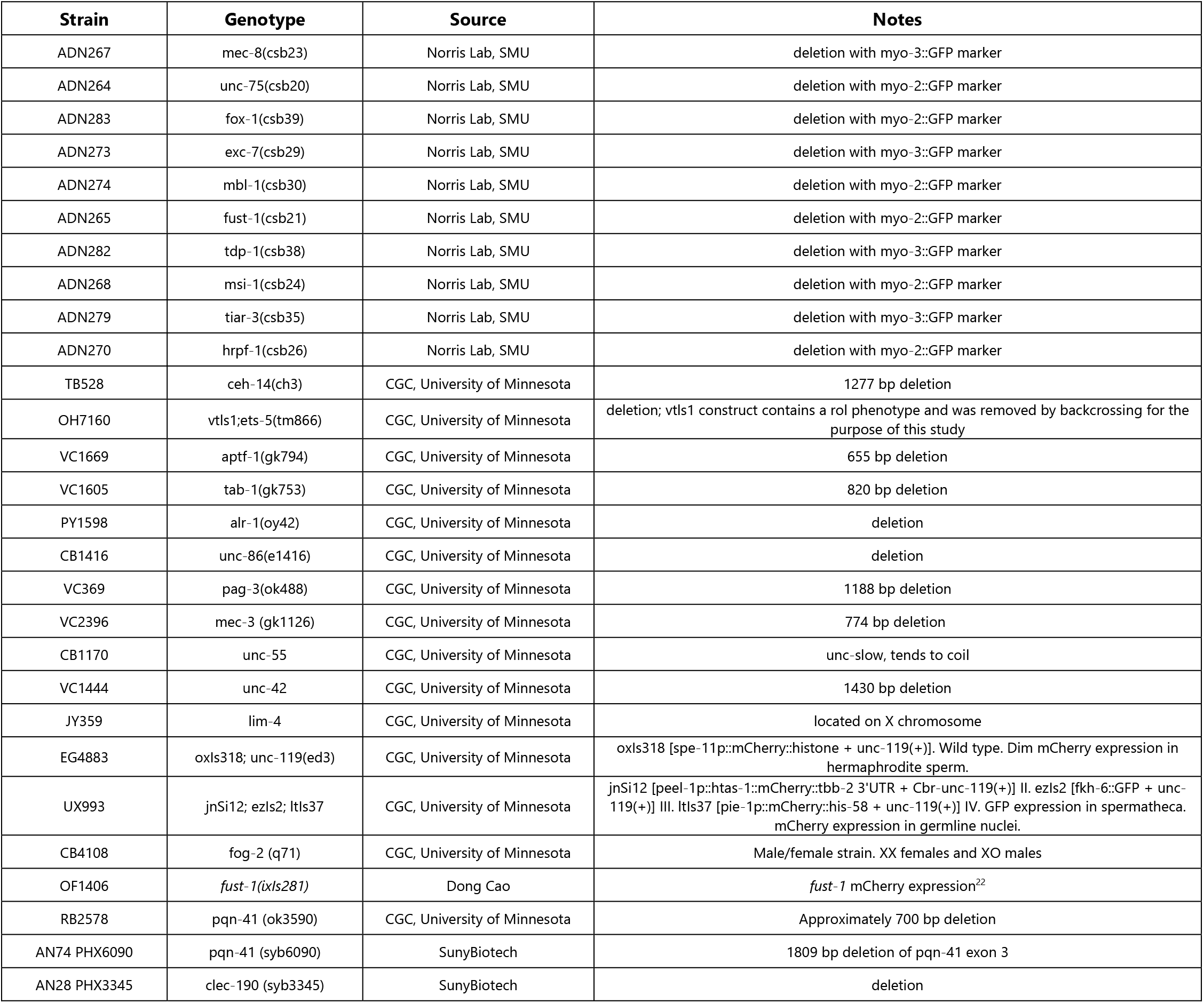

### Competitive fitness assays

Competitive fitness assays were performed as previously described to establish the relative fitness for each single mutant^20^. Briefly, four L4 larvae of each genotype were placed together on a seeded NGM plate and incubated for 5 days at 25°C. The fraction of each mutant on the plate after 5 days was calculated to generate a fitness value relative to wild-type fitness for each mutant using the formula F = (# mutant/# total)/50%. To determine double mutant fitness each double mutant was assayed with the transcription factor mutant used in the cross. Expected fitness was equivalent to the fitness of the RNA binding protein mutant in the cross. To identify double mutants with unexpected fitness, the expected fitness value for each double mutant was subtracted from its observed fitness. Synthetic fitness effects (ε) in the double mutants were calculated by F_obs_ - F_exp_. Our threshold for significance was |ε| > 0.4, and all assays were completed in triplicate.

### Larval growth assay

Worms were synchronized by standard bleaching procedure to obtain a population of L1 worms for each genotype^23^. Synchronized L1 larvae were then plated onto seeded NGM plates and cultured for 48 hours at 20°C. Developmental stages of the worms on each plate were then assessed.

### Fluorescence microscopy

Images were obtained on a Zeiss Axio Imager.Z1 microscope. Images were processed using ImageJ.

### DAPI gonad imaging

Worms were collected into microcentrifuge tubes in batches at each larval stage and at day 1 of adulthood. Whole worms were then fixed and stained with 4′,6-diamidino-2-phenylindole (DAPI). Briefly, worms were washed three times with 0.01% Tween-PBS, then frozen in 1 mL of methanol at -20°C for 5 minutes. Then, methanol was removed and worms rinsed with 1mL of 0.1% Tween-PBS. 1μL of 100ng/mL DAPI was then added to the worms in Tween-PBS, and incubated in the dark at room temperature for 5 minutes. Tubes were then rinsed with 0.1% Tween-PBS once more. Tubes were centrifuged and all solution removed from worms, then 10μL of 75% glycerol added. Worms in glycerol were then placed on agar pads on microscope slides for imaging.

### Uterine egg retention

Egg retention was measured in hermaphrodites on day 1 of adulthood. Worms were placed in 4°C refrigerator for 5-10 minutes to slow movement down, then examined under the microscope. Total eggs present in the uterus were counted.

### Lifetime egg-laying and brood size assays

Six L4 worms were placed on a seeded NGM plate and incubated at either 20°C or 25°C. The following day, and every subsequent day until egg production stopped, the 6 adults were transferred to new seeded plates. The number of progeny on each plate was counted and divided by the total number of adults to determine average egg production per day per worm. Total brood size was quantified as the sum total of eggs produced over lifespan per worm.

### Male mating efficiency

To assay male mating efficiency, four young adult males were paired with four L4 hermaphrodites, and the worms were kept at 20°C for 24 hours to allow time for mating to occur. After 24 hours the adults were removed from the plate, and progeny were given another 48 hours to develop. Progeny were then counted, and percent of male-produced cross progeny out of the total progeny were scored. Mutant males with CRISPR deletions express GFP in either the body wall muscle or the pharynx, so this fluorescence was used to identify cross progeny. For wild-type assays, males carrying myo-2::RFP which expresses bright RFP in the pharynx were used to allow scoring of cross progeny.

To measure male mating in the absence of wild-type hermaphrodite sperm, four young adult males were instead paired with four L4 *fog-2 (q71)* hermaphrodites with feminized germlines and a complete lack of sperm. For these assays, mutant males were paired with *fog-2* hermaphrodites until egg production stopped, and adults were transferred to new plates to prevent overcrowding and starvation as breeding continued. After hermaphrodites no longer continued to produce eggs, adults were removed and the total number of progeny was counted. This total was divided by the number of adult hermaphrodites present on the assay (4) to give the average brood size produced per worm. As a control, these assays were simultaneously conducted with wild-type males.

### Paired brood size assay

Four L4 wild-type males were paired with four L4 mutant hermaphrodites on a single plate, and pairs were kept paired for multiple days until egg-laying was complete. Every day, adults were transferred to new plates, and progeny left on each plate was counted to measure the total brood produced. As a control, unpaired brood assays were carried out at the same time.

### RNA sequencing and analysis

Total RNA was extracted from L4 worms using Tri reagent according to manufacturer’s protocol (Sigma Aldrich). Three biological replicates were extracted per genotype. mRNA was purified from each sample using NEBNext® Poly(A) mRNA Magnetic Isolation Module, and cDNA libraries were prepared using NEBNext® Ultra™ II RNA Library Prep Kit for Illumina, following kit protocols. Libraries were sequenced on Illumina HiSeq 2000, paired-end 150 bp reads, then mapped to the worm genome (versionWBcel235 using STAR)^24^. Gene-specific counts were tabulated for each sample using HT-Seq and statistically-significant differentially expressed transcripts were identified with DESeq2^25^. The Junction Usage Model was used to identify differentially spliced isoforms in experimental samples compared to wild type controls and quantify their expression levels by computing the ΔPSI (difference of Percent Spliced Isoform)^26^. Alternative 3’ splice site, alternative 5’ splice site, skipped “cassette” exon, and intron retention splicing events were then analyzed using custom R scripts to compare ΔPSI in fust-1;ceh-14 and tdp-1;ceh-14 with their single mutants to identify splicing dysregulation. Raw fastq files are available at the NCBI SRA (PRJNA862903).

### Reverse Transcription PCR

Relative abundances of splicing isoforms of *sav-1*, and *pqn-41* were determined by RT-PCR to confirm RNA seq results, using qScript® XLT One-Step RT-PCR Kit. The kit and reagents were used following the kit reaction protocol. Primer sequences used were as follows: *sav-1* F - GACTTCATTCAAGATCTACGG, *sav-1* R - CACTGGGAAGAGTTTGAAGCG, *pqn-41* exon 18 F - ACTACGCCTGCAACAACGTCG, *pqn-41* exon 21 R - AGCTGCTGTTGAACTTGTTGAGC

## RESULTS

### Genetic interaction screen identifies TF-RBP pairs causing synthetic fitness defects

To identify regulatory crosstalk with important functional consequences we performed a genetic interaction screen between TF and RBP mutants in *C. elegans*, focusing on evolutionarily-conserved RBPs and TFs expressed in the nervous system. To facilitate the generation of double mutants, we used existing deletion alleles which can be genotyped by simple PCR, as well as CRISPR/Cas9-mediated deletions in which we inserted a traceable fluorescent maker into the deletion locus, enabling *in vivo* monitoring of the genotype^27^ (See methods for list of genotypes used). We generated all possible double-mutant combinations of 10 RBPs and 11 TFs.

To test for genetic interactions, we measured relative fitness using a simple and quantitative competitive fitness assay^20,27^. In this assay, equal numbers of stage-matched mutant and wild-type worms are grown together on a single growth plate. The worms are given five days to develop, eat available food, and reproduce for multiple generations (Fig 1A). Then the relative proportions of mutant and wild-type worms are quantified, assigning a value to the fitness of each genotype. A mutant with an identical fitness to wild-type would grow and reproduce at the same rate as wild-type worms, resulting in a population of 50% mutants and 50% wild-type. This would yield a fitness value of 1 (Fig 1A), and increased or decreased fitness would result in values greater than or less than 1, respectively. Competitive fitness assays can identify mutants with a variety of underlying phenotypes, including lethality, developmental defects, reproductive defects, and behavioral defects^19,20^.

**Figure 1:**
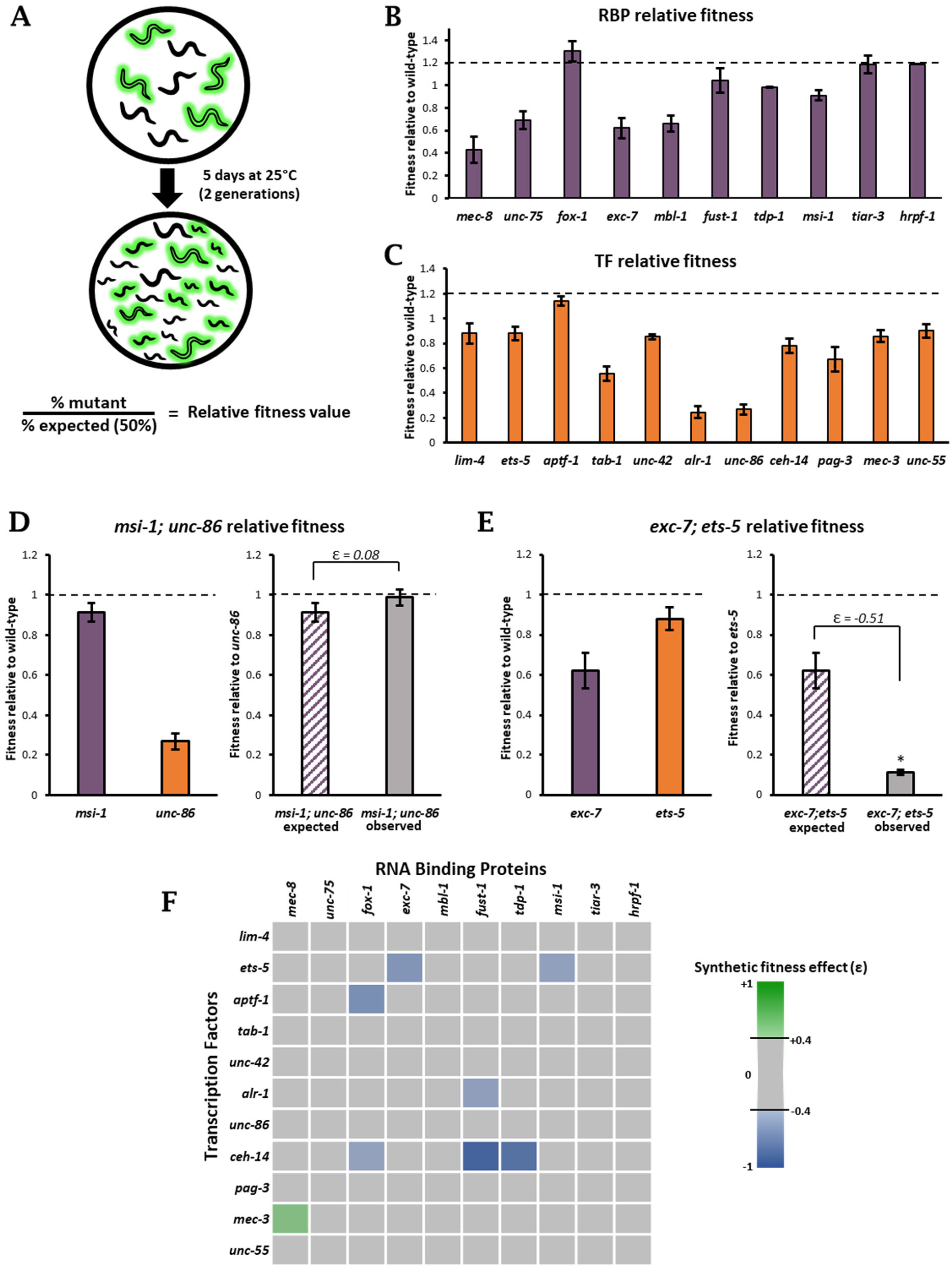
Identifying genetic interactions with competitive fitness assays. (A) Schematic for competitive fitness assays. Green worms represent fluorescent mutants, black represents wild-type. (B-C) Relative fitness values for all RNA binding protein mutants (B) transcription factor mutants (C) used in genetic interaction screen. (D-E) Double mutant relative fitness data for *msi-1; unc-86* (D) and *exc-7; ets-5* (E). Single mutants are assayed against wild-type worms and double mutants are assayed against transcription factor single mutants (F) Heat map of 110 RBP; TF double mutants created. Each square represents one double mutant. Blue squares represent synthetic negative effects on fitness (ε<-0.4), green represents synthetic positive effect (ε>0.4), and gray squares did not yield strong synthetic fitness effects (-0.4<ε<0.4)

Each RBP and TF single mutant strain was first assayed against wild-type worms to establish their respective relative fitness values (Fig 1B-C). The fitness values of the 10 RBPs and 11 TFs we assayed range from strong decreases in fitness to mild increases in fitness (Fig 1B-C). Several mutants with known behavioral defects, including *unc-86* and *mec-8*^28,29^, have significantly lower fitness than wild-type. We identified a few single mutants with novel fitness phenotypes not predicted by previously-described phenotypes, for example a reduction of fitness in *tab-1* mutants (Fig 1C). We also found that a few mutants, including *aptf-1* and *fox-1*, modestly outperform wild-type worms in our assays and have fitness values greater than 1 (Fig 1B-C)

After establishing baseline fitness values for single mutants, each RBP mutant was crossed to each TF mutant to obtain all possible RBP; TF mutant combinations, yielding a total of 110 RBP; TF double mutants. To systematically identify genetic interactions between the RBPs and TFs, we conducted competitive fitness assays in which each double mutant was competed against one of its constituent single mutants. Assuming no genetic interaction, when an RBP; TF double mutant is competed against its constituent TF mutant, the effect of the TF mutation on fitness should be equal for both the single and the double mutant. Therefore, the measured fitness of the RBP; TF should be equal to the fitness of the constituent RBP mutant when competed against wild type. For example, *msi-1; unc-86* double mutants were competed against *unc-86* mutants, and the expected fitness value was equal to the fitness of *msi-1* single mutants competed against wild type (Fig 1D). In this case, the observed fitness shows no difference from the expected fitness of the double mutant, and is therefore not considered a genetic interaction (Fig 1D).

Any deviation in the observed fitness from the expected fitness value (ε = observed – expected) constitutes a synthetic effect on fitness and signifies a genetic interaction. A positive value indicates a positive interaction, while a negative value constitutes a negative interaction. As an example, *exc-7; ets-5* double mutants were competed with *ets-5* single mutants. The expected outcome was for the measured *exc-7; ets-5* double mutant fitness value to be similar to that of *exc-7* competed against wild type, but instead *exc-7; ets-5* double mutants have significantly lower fitness than expected (Fig 1E). We set a conservative threshold of an absolute value of 0.4 or greater change in fitness (|ε| > 0.4) to be considered a strong genetic interaction. Most RBP; TF double mutants do not exhibit substantial deviations from their expected fitness value, indicating that most TFs and RBPs do not act synthetically in ways that result in changes in fitness. However, we identified eight RBP; TF double mutants with strong synthetic effects (Fig 1F).

### *aptf-1; fox-1* double mutants cause developmental delay

Performance in the competitive fitness assay depends on the ability of worms to develop, survive, reproduce, and consume food, competing for resources with other worms on the plate. Therefore, some of the underlying phenotypes that could directly impact fitness include changes in the ability to feed, move, develop and reproduce. For each double mutant that generated a significant synthetic fitness effect, where |ε|>0.4, we followed up with multiple assays to determine underlying phenotypes that contributed to a decrease or increase in fitness.

In one interesting case, we found that *apft-1; fox-1* double mutants exhibit a strong negative synthetic fitness effect, where the measured fitness is much lower than expected (Fig 2A). The RBP *fox-1* is a key regulator of *C. elegans* sex determination, while *aptf-1* is a neuronal TF important for sleep behavior^30,31^. Both factors are highly conserved, but the loss of either *fox-1* or *aptf-1* in a single mutant did not result in a measurable fitness deficit (Fig 1B-C). In the *aptf-1; fox-1* double mutant, we found moderate, but significant, reductions in egg-laying rate and pumping rate compared to *aptf-1* and *fox-1* single mutants (Fig S1). Upon further investigation, we noticed that *apft-1; fox-1* worms seemed to exhibit slower than normal growth. We measured development time from larval stage L1 to L4 and confirmed that *apft-1; fox-1* double mutants experience a significant delay in developmental timing (Fig 2B-C). 48 hours after hatching, when wild-type worms have reached the L4 larval stage, the majority of *apft-1; fox-1* double mutants are still L3 (Fig 2B). These effects are synthetic and not additive, as the constituent single mutants exhibit normal developmental timing. Therefore, although single mutants for *aptf-1* and *fox-1* display fitness equal to or greater than wild type (Fig 1B-C), when simultaneously lost they result in substantial fitness defects due to defects in developmental timing. Together, this implicates a novel role for *fox-1* and *aptf-1* in coordinating larval growth.

**Figure 2:**
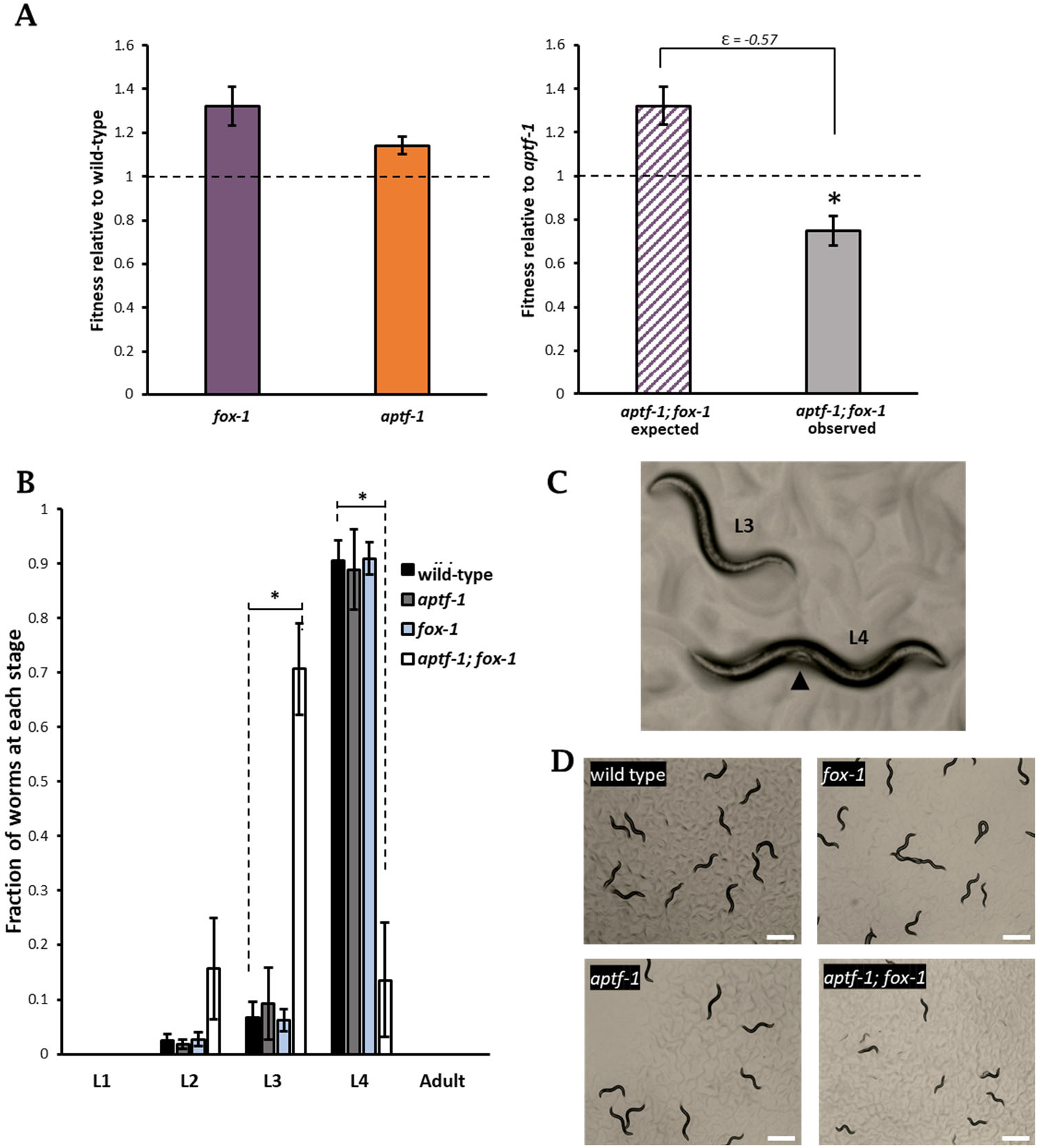
*aptf-1; fox-1* double mutants exhibit developmental delay. (A) Relative fitness data shows synthetic negative effect on fitness in *aptf-1; fox-1* double mutants. (B) Developmental timing delay in *aptf-1; fox-1* double mutants. Larval stages were assessed 48 hours after L1 worms were plated, and *aptf-1; fox-1* worms were still at L3 stage while wild-type and single mutant worms had reached L4. Asterisks indicate significant differences between *aptf-1; fox-1* and wild-type, p<0.05. (C) Representative images of *aptf-1; fox-1*, single mutants, and wild-type. Upper panel shows comparison of L3 and L4 worms. Arrow indicates characteristic white patch found in mid-body of L4, which was used to score larval stage. Lower panels show comparison of plates after 48 hours at 20°C when assays were scored.

### Reproductive defects in double mutants for TF *ceh-14* and ALS-associated RBPs *fust-1* or *tdp-1*

Two of the strongest negative synthetic effects we identified were between the homeodomain TF *ceh-14* and the RBPs *tdp-1* and *fust-1* (Fig 1F, 3A-B). *tdp-1* and *fust-1* are the *C. elegans* homologs of human TDP-43 and FUS, both implicated in ALS^32,33^. TDP-43 and FUS share many structural and functional commonalities and the roles of both RBPs in the context of disease have been well-studied^34–36^. In ALS, mutations in TDP43 or FUS cause them to mislocalize and aggregate in the cytoplasm, depleting them from the nucleus and disrupting their function^37–42^. However, their roles under non-diseased conditions are less understood.

**Figure 3:**
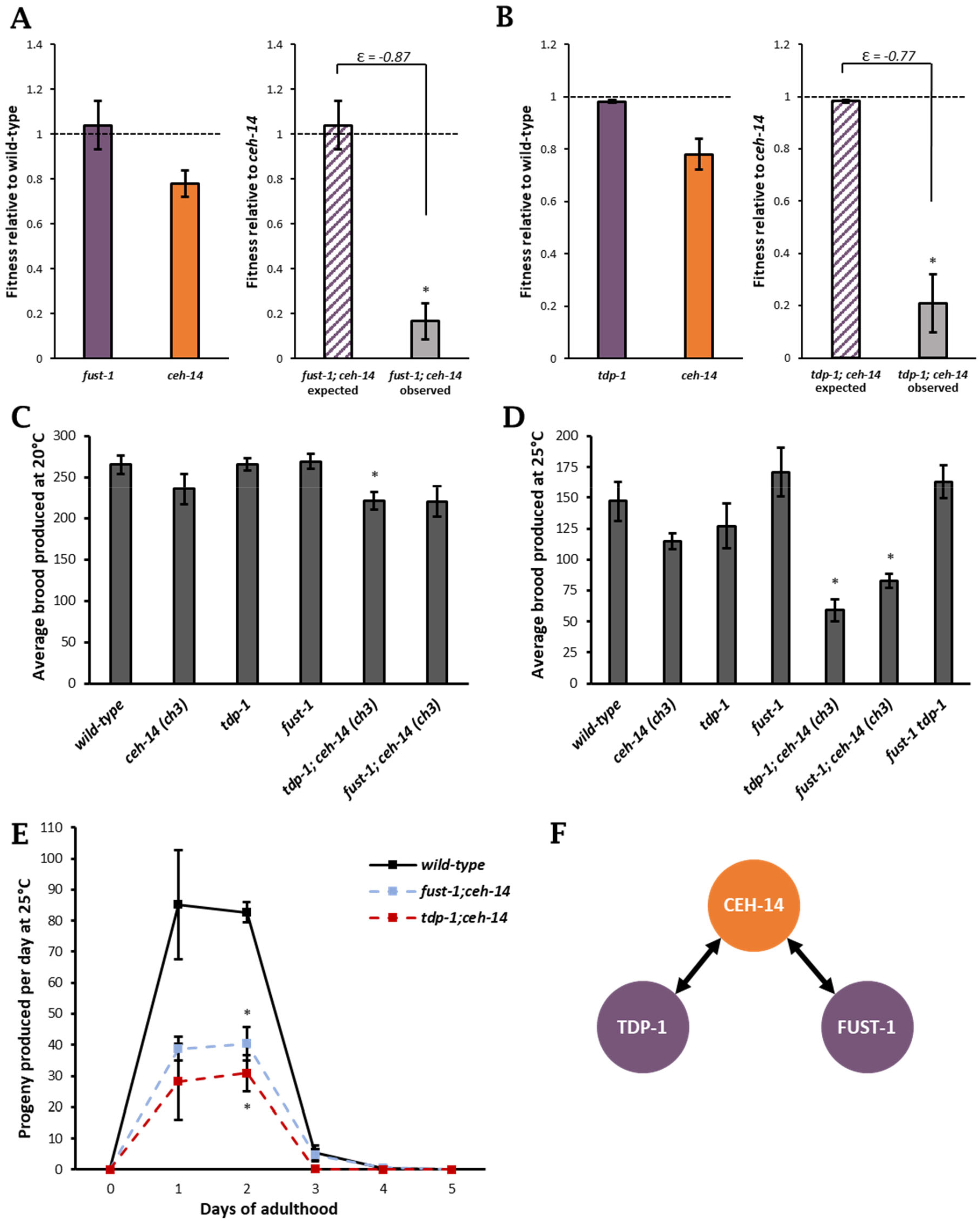
Defects in reproduction cause negative synthetic fitness effects in both *fust-1; ceh-14* and *tdp-1; ceh-14* double mutants. (A-B) Both *fust-1; ceh-14* (A) and *tdp-1; ceh-14* (B) exhibit strong negative synthetic effects on relative fitness. Asterisks indicate significant difference from expected fitness value, p <0.05 (C-D) At 20°C, differences in brood size are subtle (C), but at 25°C both *fust-1; ceh-14* and *tdp-1; ceh-14* produce significantly smaller brood sizes than wild-type, while *fust-1 tdp-1* double mutants do not exhibit a reproductive defect (D). Asterisks indicate significant difference from wild-type, p<0.05. (E) Progeny were counted every day, and brood sizes for *tdp-1; ceh-14* and *fust-1; ceh-14* are consistently lower than wild-type and single mutants. Asterisk at day 2 indicates egg production of both *fust-1; ceh-14* and *tdp-1; ceh-14* was significantly lower than that of wild-type worms, p<0.05. (F) *tdp-1* and *fust-1* exhibit negative genetic interactions with *ceh-14*, but not with each other, to affect *C. elegans* brood size.

In contrast with mammals, where loss of function of either TDP-43 or FUS is fatal^43,44^, loss of *tdp-1* or *fust-1* in *C. elegans* does not cause a strong phenotype, and indeed our *tdp-1* and *fust-1* mutants have fitness values indistinguishable from wild-type (Fig 1B). This gives us the opportunity to investigate the molecular functions of *tdp-1* and *fust-1*, taking advantage of the *ceh-14* mutant background to uncover essential roles for *tdp-1* and *fust-1* in fitness.

*ceh-14* mutants display a slight decrease in fitness compared to wild-type, but have no strong visible phenotype (Fig 1C, 3A). Only when *tdp-1* or *fust-1* mutations are combined with *ceh-14* mutations does a strong fitness defect occur (Fig 3A-B). One readily discernible commonality between *fust-1; ceh-14* and *tdp-1; ceh-14* double mutants is a reduced progeny count, based on the observation that plates of these strains appear less crowded and take longer to consume all the available food on a plate. We quantified progeny produced per worm at both 20°C and at the mildly stressful temperature 25°C^45–47^. While there is a modest decrease in brood size at 20°C, the defect becomes more pronounced at 25°C (Fig 3C-E). We measured progeny produced per day to further characterize the rate of reproduction in double mutants. We found that *tdp-1; ceh-14* and *fust-1; ceh-14* have consistently lower rates of reproduction than wild-type worms, with significantly fewer progeny produced on day 2 of adulthood compared to wild type (Fig 3E). None of the constituent single mutants exhibit this strong defect (Fig 3C-D).

To confirm that the nature of this interaction between *tdp-1* or *fust-1* and *ceh-14* is due to on-target mutations and not background effects, we generated double mutants using alternate alleles^20,48^. These new double mutants recapitulated the fertility defect, confirming the synthetic phenotypes in *tdp-1; ceh-14* and *fust-1; ceh-14* are due to on-target TF and RBP mutations (Fig S2A).

*tdp-1; ceh-14* and *fust-1; ceh-14* double mutants have fertility defects, but *tdp-1 fust-1* double mutants do not (Fig 3D). Furthermore, triple mutants *fust-1 tdp-1; ceh-14* do not have decreased fertility compared to double mutants *fust-1; ceh-14* or *tdp-1; ceh-14* (Fig S2B). Together this indicates that *tdp-1* and *fust-1* genetically interact with *ceh-14,* but not with each other, to coordinately affect reproduction in *C. elegans* (Fig. 3F). This suggests that the reproductive defects of both double mutants might stem from a shared underlying dysfunction in which the activity of both *tdp-1* and *fust-1* are required in conjunction with *ceh-14*. One plausible mechanistic scenario would be that both *tdp-1* and *fust-1* regulate a gene post-transcriptionally, while *ceh-14* regulates a second gene transcriptionally, and together both gene targets are required for fertility. In sum, we find that *tdp-1* and *fust-1*, whose human counterparts are implicated in shared disease states, also share similar physiological roles in *C. elegans*. This role in promoting fertility is only revealed in the context of the *ceh-14* mutant background.

### *fust-1; ceh-14* and *tdp-1; ceh-14* double mutants cause gonad development defects

A reduction in progeny count could be caused by a number of underlying phenotypes, including a physical inability to push eggs out of the vulva, a defect in gonadogenesis, or deficient gametes. We tested each possibility to determine the underlying causes of the double mutant phenotype. Worms with a mechanical defect in egg-laying retain eggs in the uterus^49,50^. We quantified uterine eggs in two-day-old adults and found no significant difference between double mutant and wild-type worms (Fig S3A).

We next examined the gonad to see if there were any differences in development, using DAPI (4′,6-diamidino-2-phenylindole) nuclear stain to visualize gonads within whole worms. During normal gonad development the distal tip cell (DTC) guides migration of each of the two symmetrical *C. elegans* gonad arms. The DTC guides the developing gonad out from the ventral midbody, then makes one dorsal turn, followed by a second turn towards the midbody^51,52^ (Fig 4A). This migrating tissue receives a signal to stop when the DTC crosses the midbody and reaches the vulva^53,54^. The uterine cells undergo a characteristic outgrowth, expanding and setting up the uterus centered around the vulva^55–57^.

**Figure 4:**
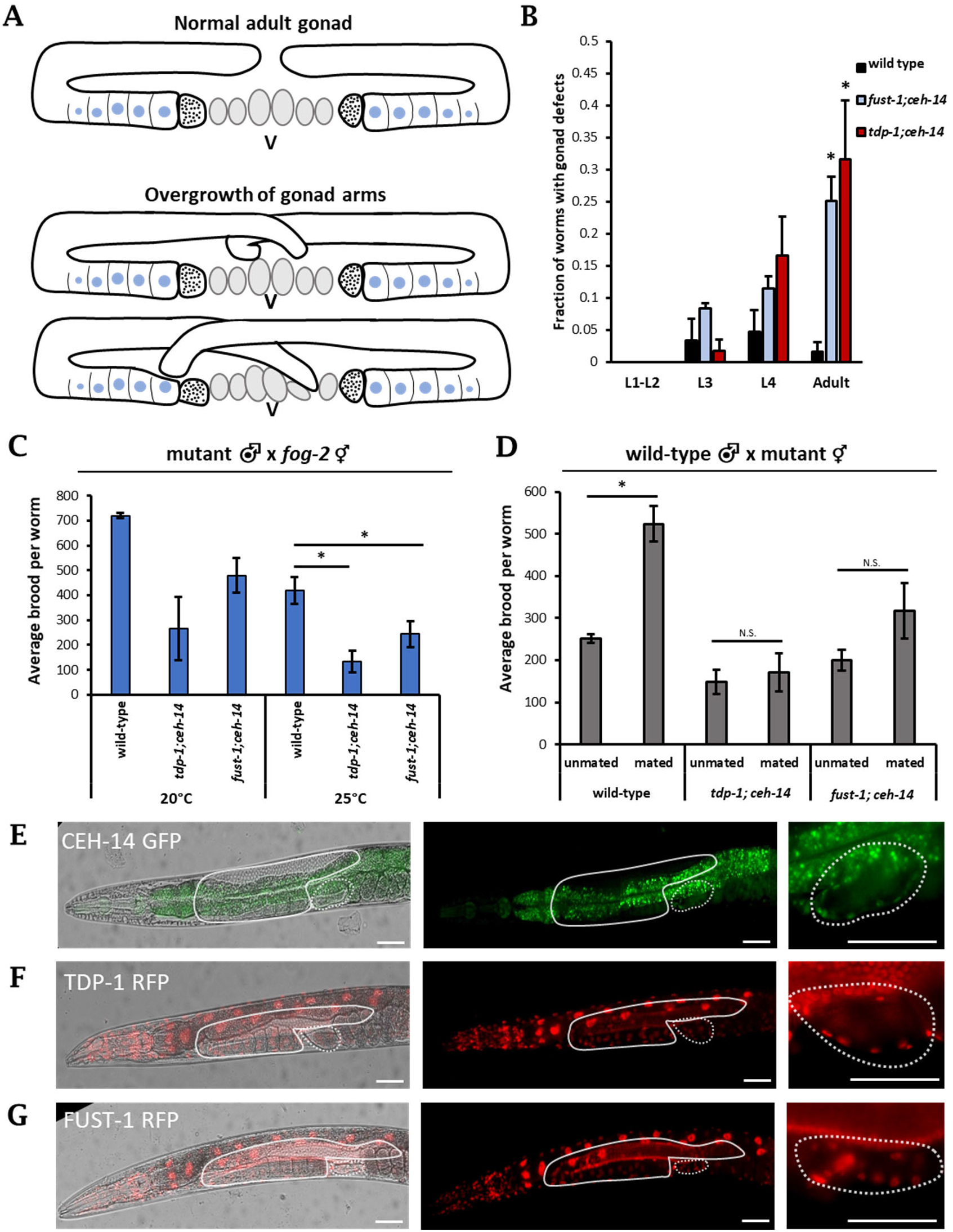
*fust-1; ceh-14* and *tdp-1; ceh-14* exhibit defects in adult hermaphrodite gonad and gametes. (A) Schematic of adult gonad in wild type compared to two examples of over-migrated gonad arms. Bottommost graphic illustrates infiltration of distal arms into both the proximal gonad and the uterus. (B) Quantification of gonad defects. Double mutants do not differ from wild-type at younger larval stages, but exhibit increased defects at adulthood. Asterisk indicates significant difference from wild-type, p<0.05. (C) Male mating with *fog-2 (q71)* feminized germline mutants. Double mutant males produced smaller brood sizes than wild type, and differences are significant at 25°C. (D) Wild-type males were mated with either wild-type or double mutant hermaphrodites. Mating with a male significantly increases brood size for wild type, but does not increase brood size in either *tdp-1; ceh-14* or *fust-1; ceh-14.* Asterisk indicates p<0.05. (E-G) Representative images show expression of CEH-14 (E), TDP-1 (F), and FUST-1 (G) in the anterior half of adult hermaphrodites. Solid outline indicates location of gonad, and dotted outline indicates location of spermatheca. Bright green visible through center of worm in (E) is gut autofluorescence. Rightmost panels show spermatheca at higher magnification. CEH-14, TDP-1, and FUST-1 are expressed in the cells making up the bag-like structure of the spermatheca. Scale bar represents 50μm

In a wild-type adult, there is typically a gap between the DTCs of the anterior and posterior arm, preventing the gonad arms from overlapping and creating ample space for the uterus (Fig 4A). When maintained at 25°C, *fust-1; ceh-14* and *tdp-1; ceh-14* double mutants present a variety of defects in gonad formation, the most common of which is apparent overmigration of the distal gonad tip in about 30% of double mutant adults (Fig 4A, S3C). The degree of overmigration varies from slight crossing of the anterior and posterior tips to more severe cases in which an overmigrated tip infiltrates the uterus or the opposite gonad arm (Fig 4A). We found no discernable differences in gonad morphology in larval stage worms, and overgrown distal arms are not observed until adulthood (Fig 4B, S3B). This suggests that rather than a dysfunction during larval gonad development, there may be a disruption in stop signaling as the distal tip reaches the vulva. Previous studies have identified similar overmigration defects in C. elegans in response to development under stress, including under conditions of changing temperature^58^, which could be related to the temperature-sensitive gonad development phenotypes observed in our double mutants.

### *fust-1; ceh-14* and *tdp-1; ceh-14* double mutants cause gamete defects

*C. elegans* hermaphrodites produce both male and female gametes and can reproduce by self-fertilization or by mating with a male. The defects we observed in self-fertilizing hermaphrodites (Fig 3) could thus stem from defects in male gametes, female gametes, or both. To determine the functionality of the male and female gametes, we conducted reciprocal mated brood assays (Fig S4). First, we mated mutant males with wild-type hermaphrodites and quantified the proportion of cross-progeny versus self-progeny to determine the efficiency of mutant male sperm. If there are no sperm defects in mutant males, the proportion of cross-progeny produced should be similar to that produced by wild-type males. At 20°C, ∼40% of progeny from wild-type male crosses are cross progeny, and at 25°C the proportion is ∼15% (Fig S4A-B). In contrast, the proportion of cross progeny from double mutant males is ∼20% at 20°C and ∼0% at 25°C. These results suggest that *tdp-1; ceh-14* and *fust-1; ceh-14* double mutant male sperm are partially defective.

In the above crosses, double mutant male sperm had to compete with wild-type sperm harbored by the self-fertile hermaphrodite. When wild-type males are crossed to hermaphrodites, male sperm is able to outcompete hermaphrodite self sperm for fertilization of oocytes^59,60^. To test whether double-mutant male sperm is fertile in the absence of competition from hermaphrodite sperm, we mated double mutant males with feminized *fog-2 (q71)* hermaphrodites which are unable to generate sperm^61^. Therefore, all progeny produced when *fog-2* hermaphrodites mate with males are cross progeny. Similar to the findings from the previous male mating assays (Fig S4), we observe a reduction in brood size for double mutant males crossed with feminized hermaphrodites, with the defect more pronounced at 25°C (Fig 4C). These findings indicate that double mutant male sperm is defective even in the absence of competition from wild-type hermaphrodite sperm.

To determine oocyte viability, wild-type males were mated with double mutant hermaphrodites. In self-fertile hermaphrodites, the total number of progeny produced is limited by the number of sperm generated. Therefore, if hermaphrodites are mated with males, the increased availability of sperm from the male will significantly increase the total progeny produced^62^. Indeed, we observe that mating with a male more than doubles wild-type brood size (Fig 4D). However, when double mutant hermaphrodites are paired with wild-type males, there is no significant increase in brood size (Fig 4D). Together these data indicate that *fust-1; ceh-14* and *tdp-1; ceh-14* double mutants are defective in both male (sperm) and female (oocyte) gametes.

### Expression of *tdp-1*, *fust-1*, and *ceh-14* overlaps in the spermatheca

To visualize where *tdp-1, fust-1,* and *ceh-14* are expressed in *C. elegans,* we endogenously tagged each gene using CRISPR/Cas9. *fust-1* and *tdp-1,* both tagged with RFP, exhibit nuclear expression in many tissues, including neurons, muscles, intestine, and the gonad. The expression of *ceh-14*, which we tagged with GFP, is more limited. As previously described, it is expressed in a subset of neurons including several in the head and tail^63^. Within the gonad, *ceh-14* exhibits nuclear expression specifically in the membrane of the spermatheca, which houses the sperm and initiates ovulation and fertilization of oocytes^64^ (Fig 4E). *fust-1* and *tdp-1* are also expressed in the nuclei of these cells (Fig 4F-G). We hypothesize that this overlapping expression in the spermatheca could be a source of their combinatorial effect on reproduction.

To investigate the development and morphology of the spermatheca in *fust-1; ceh-14* and *tdp-1; ceh-14*, we crossed each double mutant with a strain containing a spermatheca GFP reporter. Both spermathecae are present in *fust-1; ceh-14* and *tdp-1; ceh-14* adults, and they appear structurally similar to those of wild-type worms (Fig S5). We did not see any defects in the spermathecal-uterine valve or the neck of the spermatheca, both of which are indicative of spermatheca dysfunction^65–67^. This suggests that the defect in reproduction observed in our double mutants is not explained by obvious defects in spermatheca development or morphology.

### Distinct transcriptional and post-transcriptional networks in double mutants

To investigate the gene regulatory networks controlled by the three factors, we analyzed the transcriptomes of *tdp-1; ceh-14* and *fust-1; ceh-14* double mutants, as well as the constituent single mutants. At the level of gene expression, double mutants display changes in the expression of hundreds of genes compared to wild-type animals (Fig 5A, Fig S6). Such gene expression changes could be the result of (1) losing a single regulatory factor, (2) additive effects of losing both factors, or (3) synthetic/synergistic effects of losing both factors. To distinguish among these possible scenarios, we first compared gene expression changes between single mutants and double mutants. Linear regressions show that *ceh-14* accounts for the majority of gene expression changes observed in *tdp-1; ceh-14* double mutants (Fig 5B), while *tdp-1* accounts for very few gene expression changes in the double mutant (Fig 5C). Likewise, *ceh-14* accounts for the majority of gene expression changes observed in *fust-1; ceh-14* mutants (Fig S6).

**Figure 5:**
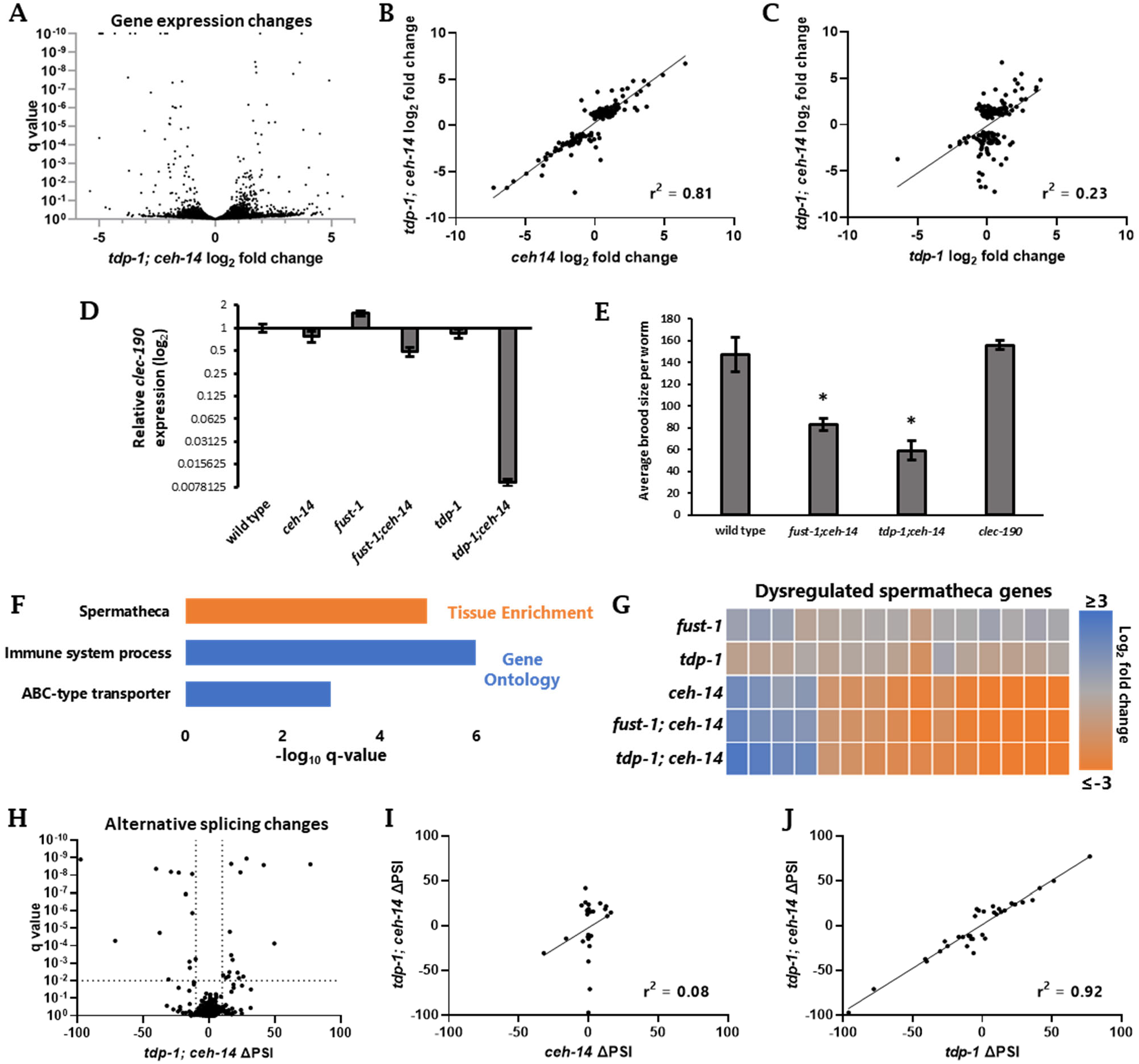
*tdp-1; ceh-14* double mutants exhibit distinct transcriptional and post-transcriptional regulation. (A) Gene expression changes of *tdp-1; ceh-14* compared to wild-type. (B-C) Linear regression showing gene expression changes for genes dysregulated (|log2fold-change| >2 and q<0.01) in *tdp-1; ceh-14* mutants, compared to *ceh-14* (B) and *tdp-1* (C) mutants. (D) qRT-PCR shows unique downregulation of *clec-190* in *tdp-1; ceh-14* double mutants. (E) *clec-190* deletion mutants do not have a fertility defect. (F) Significant tissue and gene ontology enrichment in downregulated genes of *tdp-1; ceh-14* double mutants. (G) Similar spermatheca gene dysregulation is seen in *fust-1; ceh-14* and *ceh-14* mutants. (H) Analysis of splicing changes in *tdp-1; ceh-14* compared to wild-type. (I-J) Linear regression showing splicing events dysregulated in *tdp-1; ceh-14* (|ΔPSI|>10, q<.01) compared to *ceh-14* (I) and *tdp-1* (J) mutants.

Since most gene expression changes in the double mutant are accounted for by *ceh-14* regulation, this suggests that very few gene expression changes are regulated in an additive or synergistic manner by *tdp-1* and *ceh-14*. One notable exception is the gene *clec-190*, whose expression is unchanged in single mutants, but strongly downregulated in *tdp-1; ceh-14* and modestly downregulated in *fust-1; ceh-14* mutants (Fig 5D). *clec-190* encodes a C-type lectin-like domain (CTLD) containing protein, a highly diverse protein family that fulfills a wide variety of functions^68^. Given the strong synergistic regulation of *clec-190*, we wondered whether loss of *clec-190* expression might contribute to the synergistic double mutant phenotypes. To test this, we generated a *clec-190* null mutant in which the entire coding sequence is deleted, but found that the mutant results in no discernible fertility defects (Fig 5E). Therefore, although *clec-190* represents an interesting example of combinatorial regulation, it does not on its own contribute to the double mutant phenotypes.

To test whether specific functional classes of genes are dysregulated in *tdp-1; ceh-14* double mutants, we performed Gene Ontology and Tissue Enrichment analysis. Upregulated genes have no statistically significant (q<0.01) enrichment categories, but downregulated genes are enriched in a few categories, including genes expressed in the spermatheca (Fig 5F). This is notable given the co-expression of all three factors in the spermatheca (Fig 4E-G) and the central role played by the spermatheca in fertilization. We examined all spermatheca-annotated genes with dysregulated expression in *tdp-1; ceh-14* and found that most are downregulated in the double mutant, and that *fust-1; ceh-14* double mutants have similar patterns of dysregulated gene expression (Fig 5G). Moreover, we found that *ceh-14*, but not *fust-1* or *tdp-1*, is the main driver of these changes, as *ceh-14* mutants display similar gene expression patterns to the double mutants (Fig 5G). Together these data indicate that *ceh-14* is necessary for stimulating the expression of a network of genes in the spermatheca, and motivate future work to determine whether these genes play a role in the double mutant fertility phenotypes.

Analysis of alternative splicing reveals a contrasting regulatory landscape to that of gene expression. *tdp-1* accounts for the majority of splicing changes observed in *tdp-1; ceh-14* mutants, while *ceh-14* accounts for very few splicing changes (Fig 5H-J). Likewise, *fust-1* is largely responsible for the splicing changes observed in *fust-1; ceh-14* mutants (Fig S6). As with gene expression, we observe very little additive or synergistic regulation of alternative splicing. We also observe very little overlap between genes with altered splicing regulation and genes with altered expression levels in the double mutants (Fig S6). Together, these data indicate that dysregulated genes in double mutants are either regulated transcriptionally by *ceh-14* or post-transcriptionally by *fust-1* and/or *tdp-1*.

### *tdp-1* and *fust-1* co-inhibit exon inclusion

Loss of *tdp-1 or fust-1* results in many types of dysregulated splicing, including 5’ and 3’ splice site selection, cassette exon inclusion, and intron retention (Fig 6A). One notable feature we observed is that the effect of *tdp-1* or *fust-1* mutation on cassette exons is almost exclusively an increase in exon inclusion (Fig 6B). This is in contrast with many other RNA binding proteins, which both stimulate and inhibit exon inclusion, in a context-specific manner^69^. Therefore, we conclude that *tdp-1* and *fust-1* function specifically to inhibit exon inclusion.

**Figure 6.**
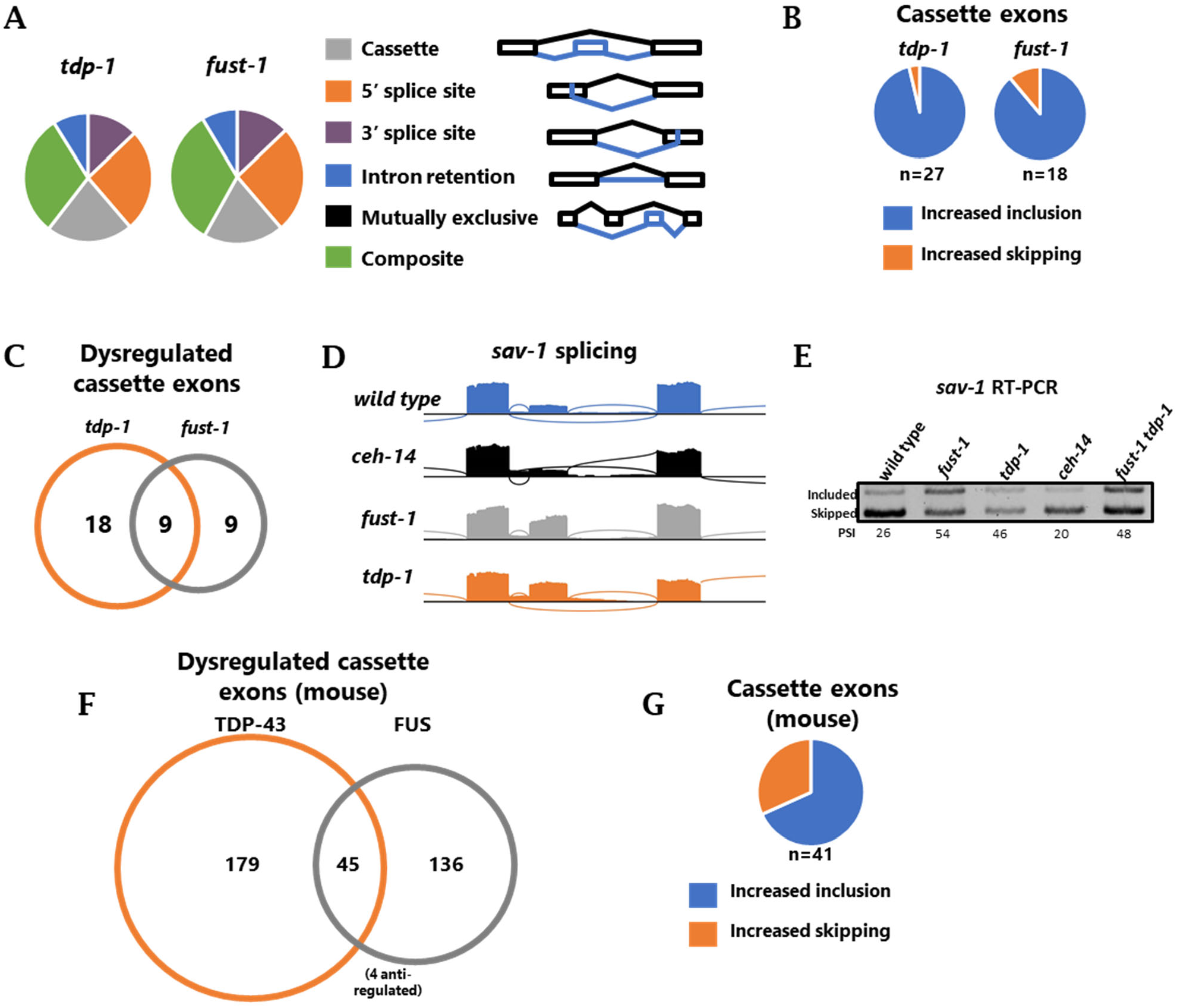
*tdp-1* and *fust-1* inhibit exon inclusion. (A) Splicing dysregulation in *tdp-1* and *fust-1* mutants (|ΔPSI|>10, q<0.01). (B) For cassette exons dysregulated in either single mutant (|ΔPSI|>10, q<0.01), increased exon inclusion in the mutant is much more common than increased skipping. (C) Overlap of cassette exon dysregulation between *tdp-1* and *fust-1* mutants (|ΔPSI|>10, q<0.01). (D-E) Increase of inclusion of *sav-1* cassette exon in both *tdp-1* and *fust-1* mutants. (E) RT PCR confirms increased exon inclusion in *tdp-1*, *fust-1*, and *fust-1 tdp-1* mutants. (F) Overlap of cassette exon dysregulation after FUS and TDP-43 knock down in mouse brain. Anti-regulated splicing events are dysregulated in both knockdowns, but in opposite directions. (G) Increased exon inclusion in majority of cassette exon dysregulation in FUS and TDP-43 mouse knock down.

We next asked whether *tdp-1* and *fust-1* inhibit expression of overlapping or distinct alternative exons. Strikingly, we found that half of *fust-1*-regulated cassette exons are also regulated by *tdp-1* (Fig 6C), and that the direction of splicing change is always concordant, with increased exon inclusion in both mutants. An example of an exon repressed by both factors is in the gene *sav-1*, which harbors an unannotated cassette exon. In wild-type or *ceh-14* mutants, this exon is predominantly skipped, but in either *fust-1* or *tdp-1* mutants, the exon becomes predominantly included (Fig 6D-E). In *fust-1 tdp-1* double mutants, the exon is included at levels similar to either single mutant (Fig 6E), suggesting that *tdp-1* and *fust-1* do not act synergistically, but rather are both simultaneously required to repress *sav-1* exon inclusion.

Given the striking concordance of inhibition of exon inclusion by *fust-1* and *tdp-1*, we next asked whether such activity is an evolutionarily-conserved attribute of the two RBPs. This would be of particular interest given both factors’ prominent links to the human neuronal disorders ALS and FTD^70^. To this end we re-analyzed data in which either of the mouse homologues (FUS or TDP-43) was knocked down in mouse brains and splicing analyzed by microarray^42^. Focusing on cassette exons, we found a substantial overlap between the regulatory activity of FUS and TDP-43 (Fig 6F). As in our *C. elegans* experiments, exons co-regulated by both FUS and TDP-43 in mouse brain tend to be inhibited by both factors (Fig 6G). Thus, *tdp-1/TDP43* and *fust-1/FUS* have a propensity to coordinately inhibit exon inclusion both in *C. elegans* and in mouse brain.

### Aberrant exon inclusion of *pqn-41* contributes to fertility defects

One intriguing example in which *C. elegans tdp-1* (but not *fust-1*) inhibits exon inclusion is found in *pqn-41*, a gene encoding a polyglutamine-containing protein. This gene harbors an alternative cassette exon which in wild-type conditions is primarily skipped, but in *tdp-1* mutants is primarily included (Fig 7A-C). *pqn-41* was previously shown to be important for proper developmental cell death of the linker cell in the gonad of male *C. elegans*^71^, which prompted us to ask whether *pqn-41* might contribute to the gonad development or fertility defects observed in *tdp-1; ceh-14* mutant hermaphrodites. We obtained a potentially null deletion allele, *pqn-41(ok3590)*^72^, and tested fertility. We found that brood sizes of *pqn-41* mutants are significantly lower than wild-type worms, and that these fertility defects are particularly pronounced at higher temperatures (Fig 7D). Therefore, loss-of-function *pqn-41* mutants have similar temperature-sensitive fertility defects to *tdp-1; ceh-14*.

**Figure 7.**
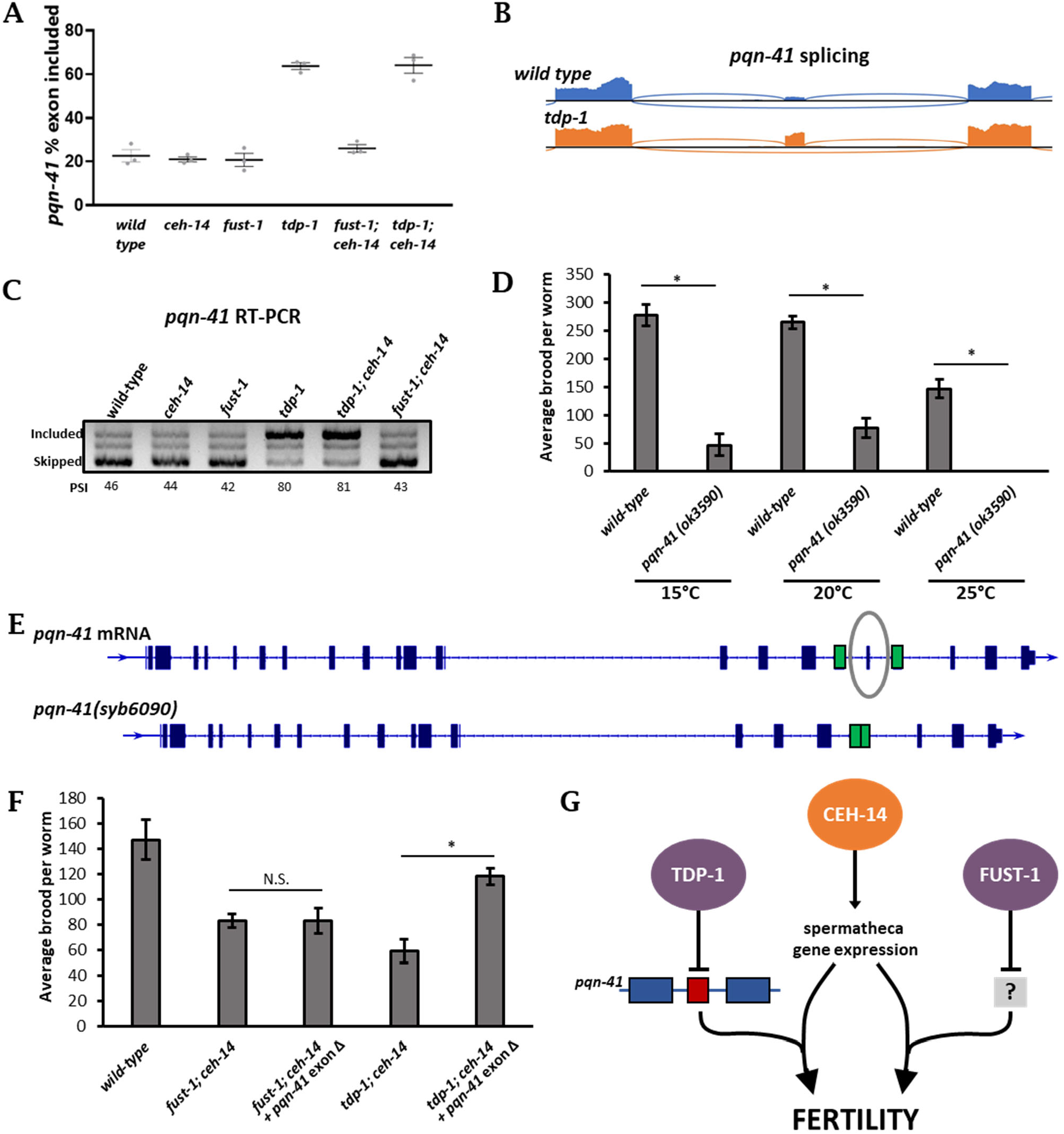
*tdp-1* inhibition of *pqn-41* alternative exon splicing contributes to fertility. (A-C) Increased inclusion of *pqn-41* cassette exon in *tdp-1* and *tdp-1; ceh-14* mutants. (D) Reduced fertility in *pqn-41 (ok3590)* deletion mutants. Asterisks indicate significant difference from wild-type, p<0.05. (E) mRNA sequence of *pqn-41*, with alternative exon circled. CRISPR/Cas9 was used to delete the exon with surrounding introns, and flanking exons (shown in green) were fused^73^ (F) Deletion of *pqn-41* exon rescues brood size in *tdp-1; ceh-14* double mutants. Asterisk indicates significant difference from *tdp-1; ceh-14* brood size, p<0.05. (G) Molecular model of TDP-1 – CEH-14 and FUST-1 – CEH-14 interactions affecting *C. elegans* fertility.

We next generated a *pqn-41* mutant using CRISPR/Cas9 in which the alternative exon is removed and the flanking exons are precisely fused together, thereby forcing expression of the exon skipped version (Fig 7E). We then crossed this *pqn-41* exon-deletion mutant into a *tdp-1; ceh-14* background, thus restoring *pqn-41* to the isoform most abundant under wild-type conditions (exon skipped). Remarkably, these triple mutants (*tdp-1; pqn-41[exon-skipped]; ceh-14*) exhibit a strong rescue of brood size at 25°C compared to *tdp-1; ceh-14* double mutants (Fig 7F). This suggests that aberrant *pqn-41* exon inclusion plays a major role in the fertility defects observed in *tdp-1; ceh-14* double mutants.

In contrast with *tdp-1; ceh-14* double mutants, *fust-1; ceh-14* double mutants do not exhibit *pqn-41* splicing defects. Likewise, crossing the *pqn-41[exon-skipped]* into the *fust-1; ceh-14* double mutant does not cause an increase in brood size (Fig 7F), suggesting that the rescue of *tdp-1; ceh-14* by *pqn-41[exon-skipped]* is mechanistically linked to the mis-splicing of *pqn-41* caused by *tdp-1* loss of function. Together these results highlight a new role for the polyglutamine gene *pqn-41* in fertility, and indicate that *pqn-41* mis-splicing is a major cause of the fertility defects observed in *tdp-1; ceh-14* double mutants.

In sum, using a systematic combinatorial genetic interaction screen, we found that two RBPs, *fust-1* and *tdp-1*, are both required in the context of a *ceh-14* mutant background to maintain fitness and fertility in *C. elegans*. These two RBPs have both been implicated in ALS and FTD in humans, and we now identify a common physiological role for both RBPs in *C. elegans*. Both RBPs have overlapping roles in inhibiting exon inclusion, pointing to shared molecular activities, and a potential molecular basis for the physiological roles for the two RBPs described here. Failure to inhibit exon inclusion in the *pqn-41* gene is a major cause of the fertility defects in *tdp-1; ceh-14* double mutants, thus providing a mechanistic link between the molecular activity of the TDP-1 RBP and the fertility phenotype observed in *tdp-1; ceh-14* double mutants (Fig 7G).

## DISCUSSION

### Novel genetic interactions across gene regulatory layers

We took a systematic genetic interaction approach to identify cross-regulatory genetic interactions in which a TF and RBP are combinatorially required for phenotypes affecting organismal fitness. This screen revealed a number of TF-RBP pairs required for phenotypes including fitness, development, and fertility. The strongest of these genetic interactions involves the homeodomain TF *ceh-14* and either of the ALS-associated RBPs *tdp-1* or *fust-1*.

Transcriptome analysis revealed distinct regulatory networks, in which *ceh-14* regulates transcription, and *tdp-1/fust-1* regulate splicing, with few genes additively or synergistically regulated, and few genes regulated at both transcriptional and post-transcriptional levels. Therefore, it seems likely that the synthetic fertility phenotypes result from the combination of distinct gene dysregulation events. We identify one such dysregulation in the alternatively-spliced exon in *pqn-41*. Mis-splicing of this exon is a major contributor to the phenotype, as restoring exon skipping rescues fertility defects of *tdp-1; ceh-14* double mutants. Mis-splicing of this exon in isolation does not cause fertility defects, as *tdp-1* single mutants mis-splice *pqn-41* at the same levels as *tdp-1; ceh-14* mutants, but do not have fertility defects. We therefore speculate that mis-splicing of *pqn-41*, in combination with altered expression of one or more *ceh-14* target genes, results in fertility defects (Fig 7G). A number of genes with spermatheca expression dependent on *ceh-14* (Fig 5G) represent promising candidates.

These results highlight the utility of the systematic reverse-genetic interaction approach both for understanding relationships between regulatory factors and for understanding the roles of individual factors whose regulatory roles are only apparent when redundant or compensatory pathways are simultaneously perturbed^16^. In this case, a shared molecular and physiological role for *tdp-1* and *fust-1* is revealed by their shared genetic interaction profile. Future studies characterizing additional genetic interactions identified here may shed light on novel physiological roles for additional RBPs and TFs.

### *fust-1* and *tdp-1* interact with *ceh-14* to affect *C. elegans* fertility

The hermaphrodite spermatheca is a key site of overlapping expression for *tdp-1*, *fust-1*, and *ceh-14*, and double mutants have reduced sperm efficiency. Signaling from somatic gonad cells such as spermathecal cells is required for germline development and function, as ablation of specific spermatheca cells results in defective germ cell function, and even sterility^74^. We hypothesize that faulty signaling between spermathecal cells and germ cells might explain the defects in fertility and gonad development observed in our double mutants. The double-mutant fertility phenotype is particularly pronounced at 25°C, which is a mildly stressful temperature for wild-type worms, causing modest defects in fertility and gonad development^23,47,58^. Temperatures higher than 25°C result in damage to sperm and strong fertility defects^75,76^. We speculate that *tdp-1; ceh-14* and *fust-1; ceh-14* double mutants are deficient in their heat stress responses^45,46^, and are therefore unable to maintain normal homeostasis under mild heat stress. Therefore, temperatures that cause mild fertility defects in wild-type animals result in strong defects in double mutants.

How might *pqn-41* splicing contribute to this temperature-specific fertility defect? The PQN-41 protein is abundant with glutamines, and contains a particularly polyglutamine-rich domain at the C-terminus^71^ . This domain begins immediately downstream of the alternative exon (Fig S7). We hypothesize that, like other polyglutamine proteins such as the Huntington protein, PQN-41 is subject to pathogenic aggregation^77^. If the exon-included isoform of PQN-41 is particularly prone to aggregation, and if stressful conditions such as higher temperatures further increase the likelihood of aggregation, this could lead to temperature-sensitive defects. Indeed, there is evidence that PQN-41 forms aggregates *in vivo*^71^, but a full mechanistic test of this hypothesis awaits further investigation. It will be interesting in future studies to investigate whether there is a related prion-like polyQ protein underlying the similar fertility defects of *fust-1; ceh-14* double mutants.

### *fust-1* and *tdp-1* co-inhibit exon inclusion

Identification of shared phenotypes between *tdp-1; ceh-14* and *fust-1; ceh-14* double mutants led to the observation that *tdp-1* and *fust-1* also have shared effects on the transcriptome. Most notably, they both act to inhibit inclusion of alternatively-spliced cassette exons, including many inhibited by both RBPs. *fust-1* and *tdp-1* do not appear to act redundantly, as *fust-1 tdp-1* double mutants do not result in increased exon inclusion compared to either of the single mutants. Rather, *fust-1* and *tdp-1* are both simultaneously required for inhibition of these exons. One plausible explanation for this finding is that both RBPs bind together to specific pre-mRNAs where they act in concert to prevent aberrant exon inclusion.

The activity of *C. elegans tdp-1* and *fust-1* in inhibiting exon inclusion is also a shared feature of mammalian TDP-43 and FUS. We find that knockdown of TDP-43 or FUS in mouse brain^42^ results in aberrant exon inclusion, and that many of these exons are co-inhibited by both TDP-43 and FUS. This is interesting in light of recent findings suggesting a pathologically-relevant role for TDP-43 in inhibiting cryptic exons. Exons are sometimes classified as cryptic if they exhibit low inclusion levels, lack of evolutionary conservation, and/or propensity to disrupt the function of the gene they reside in^78,79^. TDP-43 has been identified as an inhibitor of cryptic exons^78,79^, and recent evidence implicates aberrant inclusion of two different cryptic exons in the genes *STMN2* and *UNC13A* as potential causative mechanisms underlying TDP-43 pathology in ALS^80–83^.

Our findings are consistent with a role for *tpd-1*/*TDP-43* in inhibiting aberrant exon inclusion, and we extend this observation to also include a role for *fust-1*/*FUS* in inhibiting exon inclusion. This leads us to speculate whether FUS-related pathogenesis might also be mechanistically linked to inappropriate inclusion of exons inhibited by FUS. Previous work on mammalian TDP-43 and FUS has concluded that the two RBPs share many common RNA targets, but also have considerable non-overlapping regulatory functions^42,84^. We focused here on the regulation of cassette exons, and found substantial overlap between the RBPs in inhibiting exon inclusion. It will therefore be interesting to ask whether aberrant exon inclusion underlies FUS-mediated pathology in an analogous way to that of TDP-43-mediated pathology.

Many of the exons identified as targets of *tdp-1* and/or *fust-1* in *C. elegans* have attributes of cryptic exons as well. For example, the alternative exons in *sav-1* and *pqn-41* are expressed at low levels in wild-type (Fig 6D, 7B), and in the case of *sav-1* the exon is unannotated. In the case of *pqn-41*, failure of *tdp-1* to inhibit exon inclusion leads to detrimental phenotypes (fitness and fertility defects). This is an interesting parallel to the pathogenic consequences of TDP-43 failing to inhibit cryptic exon inclusion in the *STMN2* or *UNC13A* genes, and suggests that inhibition of aberrant exon inclusion may be an evolutionarily-conserved feature of *tdp-1/TDP-43* and *fust-1/FUS*.

## ACKNOWLEDGEMENTS

Thank you to the entire Norris Lab for ideas, suggestions, and critical reading of the manuscript. Funding was provided by: National Institute of General Medical Sciences of the National Institutes of Health [R35GM133461]; National Institute of Neurological Disorders and Stroke of the National Institutes of Health [R01NS111055]. Author contributions-Conception/design, drafting/revision, performance of experiments: MT, OM, AN.

## SUPPLEMENTAL FIGURES

**Supplemental figure 1.**
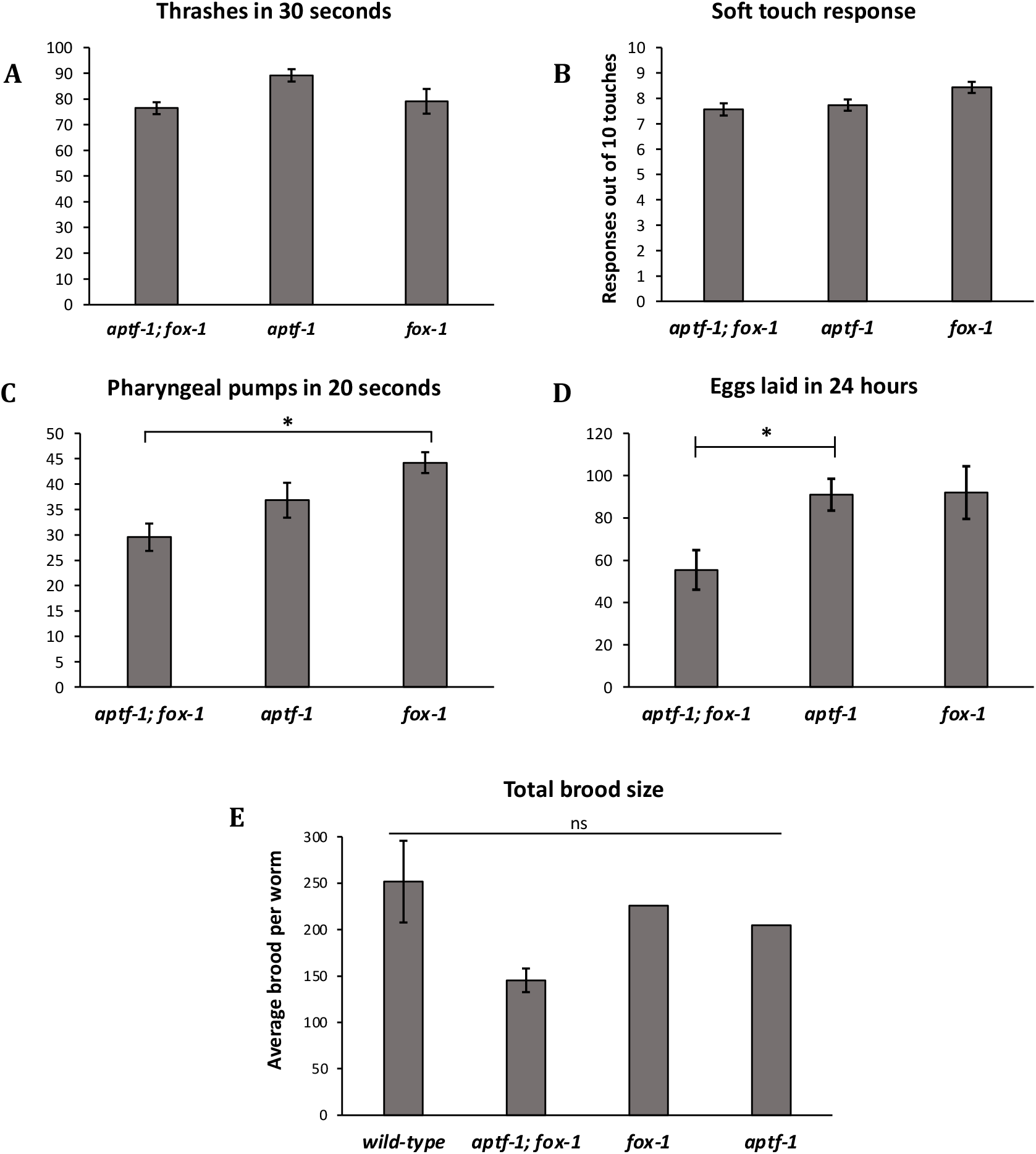
*aptf-1; fox-1* double mutants exhibit slowed rates of egg laying and pharyngeal pumping compared to single mutants. (A) Number of thrashes per L4 worm in 30 seconds. No significant differences were observed between groups. (B) Soft touch assays were performed with an eyelash hair pick. No significant differences were observed. (C) Number of pharyngeal pumps in 20 seconds was counted. Asterisk indicates a significant reduction in pumps in *aptf-1; fox-1* compared to *fox-1* single mutants, p < 0.05. (D) Early adult worms were individually plated and given 24 hours to lay eggs. Asterisk indicates significantly reduced eggs were counted on *aptf-1; fox-1* plates compared to *aptf-1,* p < 0.05. (E) Total brood size produced over the adult lifespan of each worm was quantified. No significant difference was observed between groups.

**Supplemental Figure 2.**
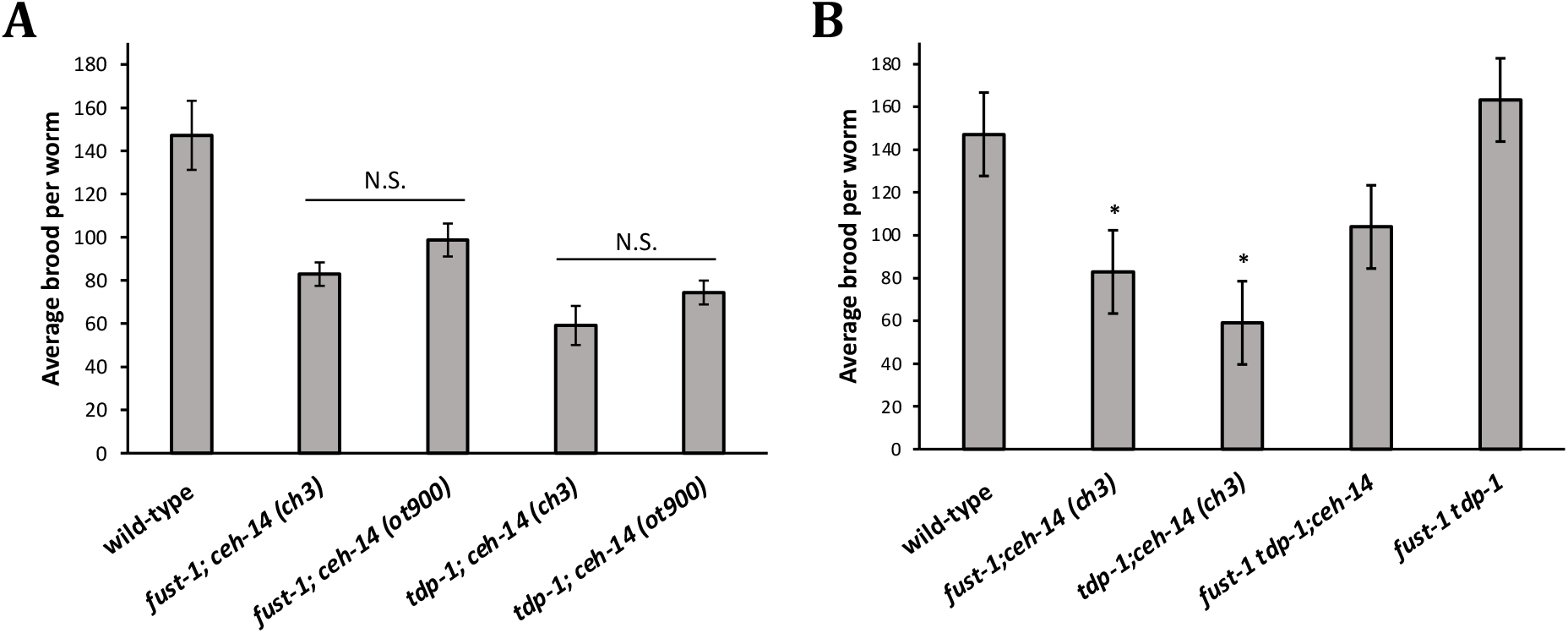
f*u*st*-1; ceh-14* and *tdp-1; ceh-14* phenotypes are recapitulated with alternative alleles. (A) Both double mutants were generated using *ceh-14 (ot900)*, which contains a full deletion of *ceh-14*. No significant differences in brood size were measured between double mutants generated with the original *ceh-14 (ch3)* allele and those generated with *ceh-14 (ot900)*. (B) A *fust-1 tdp-1; ceh-14* triple mutant did not exhibit a worsened brood size defect compared to *fust-1; ceh-14* and *tdp-1; ceh-14* double mutants. Asterisk indicates significant difference from wild-type, p<0.05. *fust-1 tdp-1; ceh-14* brood size is not significantly different from that of wild-type.

**Supplemental Figure 3.**
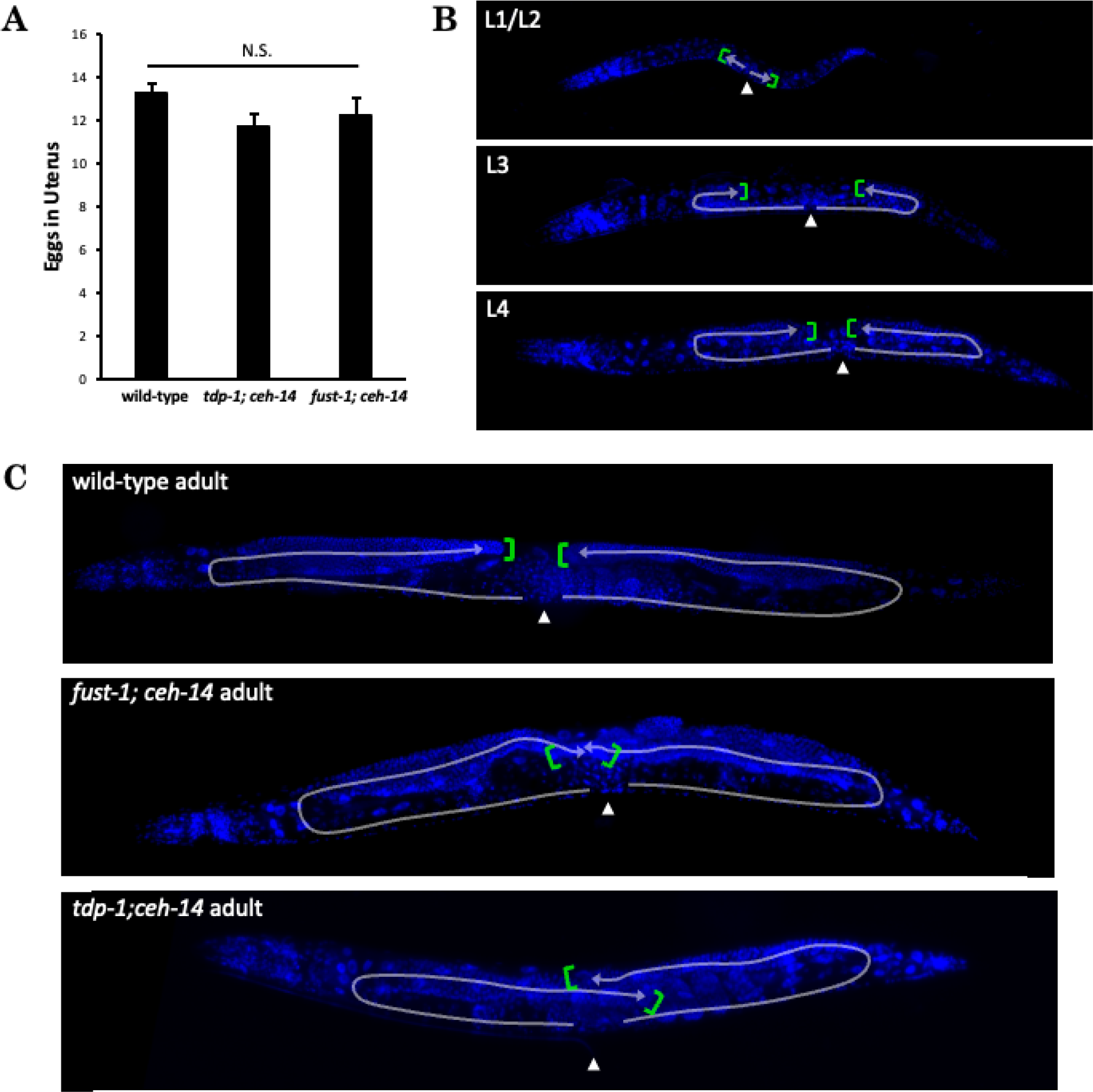
f*u*st*-1; ceh-14* and *tdp-1; ceh-14* exhibit defects in adult hermaphrodite gonad. (A) Double mutants do not exhibit egg retention. (B) Representative images of DAPI-stained larval hermaphrodites. (C) Representative images of DAPI-stained wild-type and double mutant adult hermaphrodites. *fust-1; ceh-14* and *tdp-1; ceh-14* exhibit overlapping distal tips. Arrows indicate path of gonad development. Arrow heads denote location at midbody where vulva is present in adult. Green brackets indicate approximate location of distal tip.

**Supplemental Figure 4.**
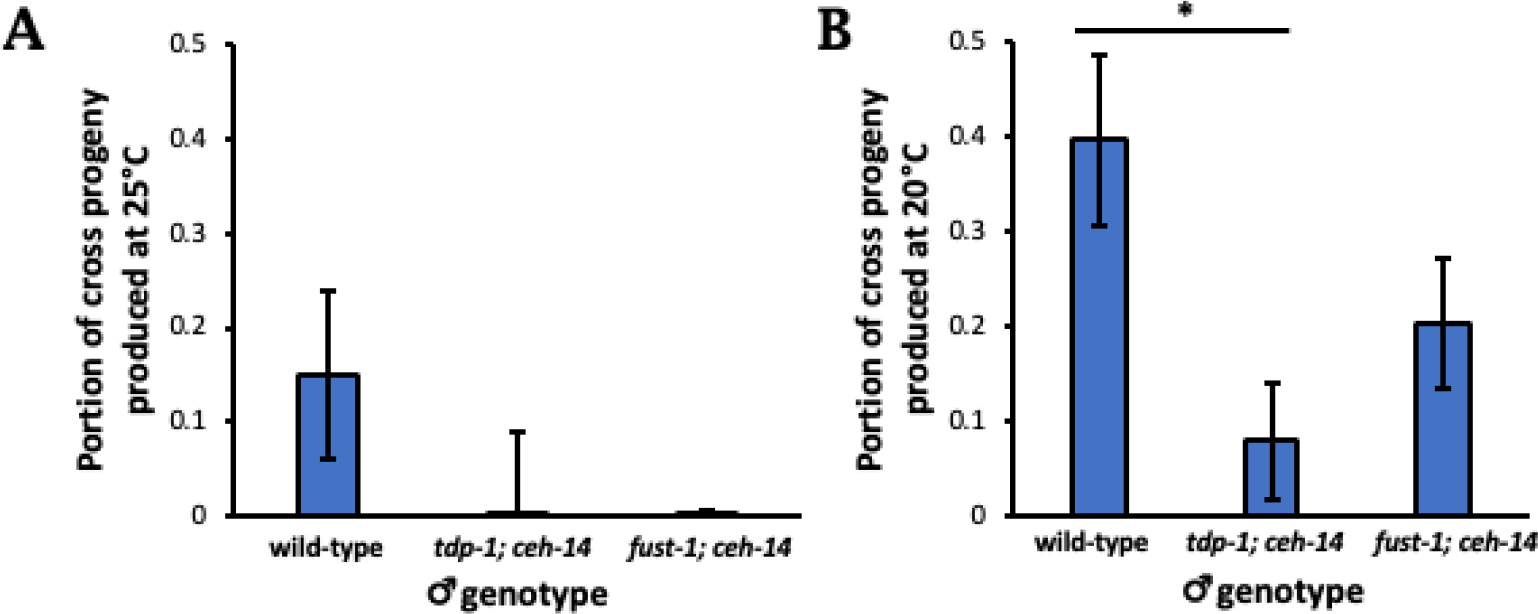
Male mating defects in double mutants. *tdp-1; ceh-14* and *fust-1; ceh-14* males maintained at 25°C (A) and at 20°C (B) were paired with wild-type hermaphrodites, and production of cross-progeny was measured. Asterisk indicates significant difference from wild-type cross progeny, p<0.05.

**Supplemental Figure 5.**
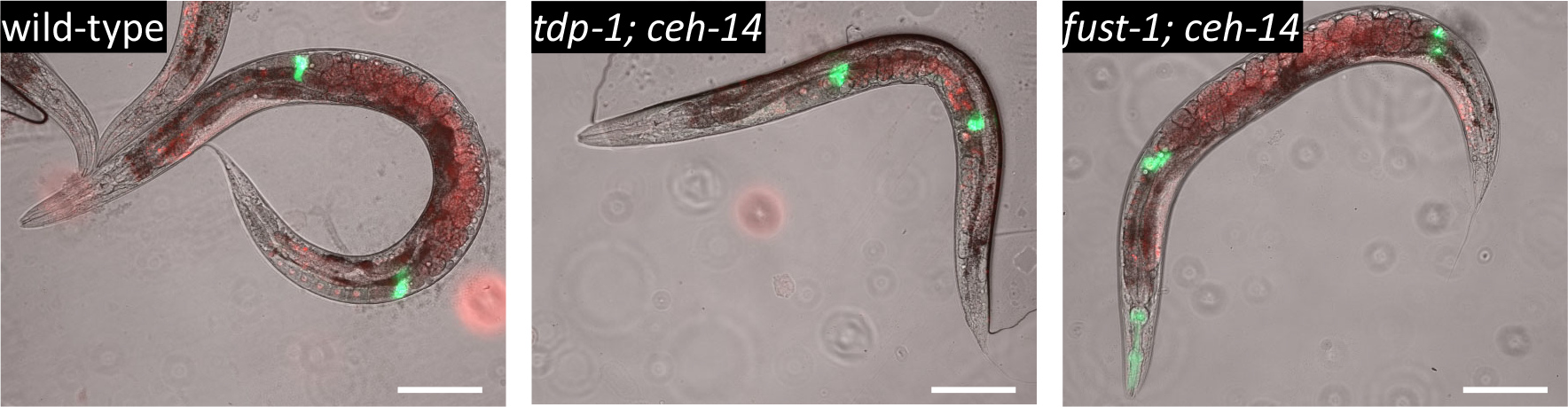
Spermatheca of *tdp-1; ceh-14* and *fust-1; ceh-14* double mutants are morphologically similar to wild-type. UX993 worms containing spermatheca GFP and germline RFP were crossed into *tdp-1; ceh-14* and *fust-1; ceh-14* double mutants to visualize spermatheca morphology. No differences in morphology or development of spermatheca were detected in either double mutant. Representative images show day 1 adults of wild-type and double mutants containing the UX993 transgenes.

**Supplemental Figure 6.**
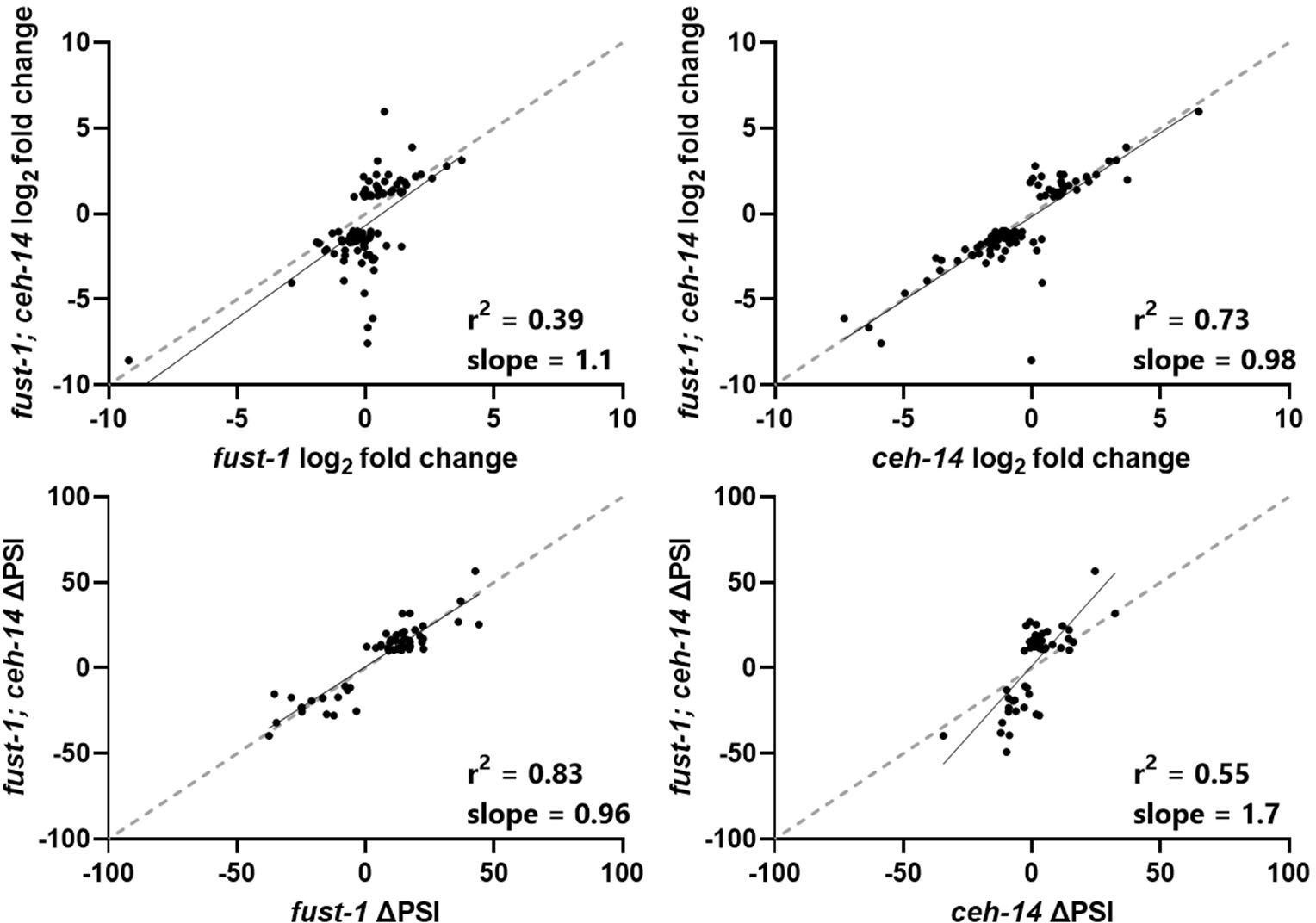
f*u*st*-1; ceh-14* exhibit distinct transcriptional and post-transcriptional regulation. Linear regressions showing gene expression or splicing changes for genes dysregulated (for gene expression|log2fold-change| >2 and q<0.01, and for alternative splicing |ΔPSI|>10, q<.01) in *fust-1; ceh-14* mutants, compared to *ceh-14* and *fust-1* mutants.

**Supplemental Figure 7.**
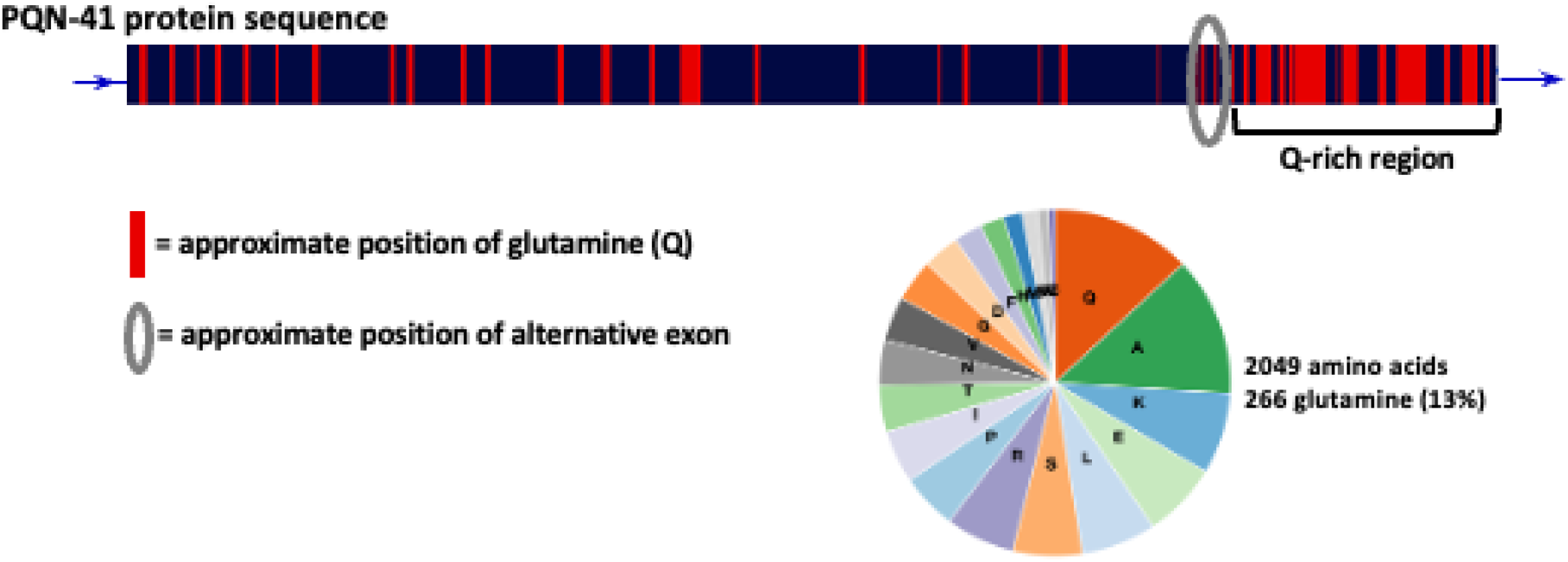
p*q*n*-41* amino acid sequence. Approximate locations of glutamine are highlighted in red, and Q-rich region is shown downstream of the circled alternative exon. pqn-41 amino acid composition reveals a bias towards glutamine^86^.

